# Species-dependent actions of the Goαb selective adenosine A_1_ receptor agonist BnOCPA

**DOI:** 10.1101/2022.12.02.518704

**Authors:** Emily Hill, Xianglin Huang, Ivana Del Popolo, Circe La Mache, Martin Lochner, Graham Ladds, Bruno G. Frenguelli, Mark J. Wall

**Affiliations:** School of Life Sciences, University of Warwick, Coventry, CV4 7AL; Department of Pharmacology, University of Cambridge, Tennis Court Road, Cambridge, CB2 1PD; Institute of Biochemistry and Molecular Medicine, University of Bern, 3012 Bern, Switzerland

## Abstract

We have previously reported that in rat hippocampal area CA1, the A_1_R-selective agonist, BnOCPA, potently inhibited excitatory synaptic transmission but did not cause membrane hyperpolarisation in CA1 pyramidal neurons, as would be expected of A_1_R agonists. This functional discrimination by BnOCPA may arise from its ability, in cAMP inhibition assays, to selectively activate only Gob out of the six Gαi/o subtypes. This may explain why BnOCPA is a potent analgesic that does not cause sedation or cardiorespiratory depression in the rat. Since many preclinical studies are performed using mice, we have here investigated whether BnOCPA’s functional discrimination extends to the mouse. While the potency of BnOCPA against the inhibition of hippocampal synaptic transmission was comparable between rats and mice, we discovered that low concentrations of BnOCPA hyperpolarised mouse CA1 neurons and reduced both their input resistance and firing rate in an A_1_R-dependent manner. In interleaved experiments we confirmed our previous observations in the rat that concentrations of BnOCPA equivalent to those tested in the mouse had little or no effect on membrane potential or input resistance. Using NanoBRET binding we established that BnOCPA had similar affinity at the mouse and rat A_1_Rs, and displayed little difference in G protein coupling, as determined using the TRUPATH assay. Thus, although the mechanism for the loss of BnOCPA functional selectivity between pre- and postsynaptic receptors in the mouse hippocampus is currently unclear, it may stem from differences in expression of the individual G proteins subunits or the coupling to murine K^+^ channels.

**Short summary:** We describe the differential actions of the selective A_1_R agonist BnOCPA in mouse vs rat hippocampus. In mice, BnOCPA does not show a selectivity between pre and postsynaptic A_1_Rs, unlike in rats. This may stem from differences in the G protein coupling to K^+^ channels.

## Introduction

The purine nucleoside adenosine plays a vital role in many physiological and pathological processes across all mammalian organ systems (Borea et al., 2018). These actions, mediated by four G protein-coupled receptors (GPCRs) include regulation of neuronal excitability, control of cardiac contractility and modulation of vascular tone. Of the four adenosine GPCRs, the adenosine A_1_ receptor (A_1_R) has attracted considerable interest due to its powerful inhibitory influences on neuronal and cardiovascular function. In particular, the endogenous agonist adenosine inhibits excitatory glutamatergic synaptic transmission and hyperpolarises postsynaptic neurones, actions that are mimicked by conventional A_1_R agonists such as N6-cyclopentyladenosine (CPA) and 5’-N-Ethycarboxamidoadenosine (NECA).

In screening A_1_R agonists derived from a search for molecules suitable as fluorescent ligands (Knight et al., 2016), we previously utilised molecular dynamics (MD) simulations and Gαi/o subunit- and β-arrestin-specific cellular signalling assays to make the unprecedented observation that one A_1_R-selective agonist, benzyloxy-cyclopentyladenosine (BnOCPA), exclusively activates Gob among the six members of the Gαi/o family of G protein subunits (Wall et al., 2022). Moreover, through a combination of CNS electrophysiology, physiological recordings of cardiorespiratory parameters, and the use of a clinically-relevant model of chronic neuropathic pain, we demonstrated selective activation of native A_1_Rs and the delivery of potent analgesia without the sedation, motor impairment, bradycardia, hypotension or respiratory depression normally associated with conventional A_1_R agonists (Wall et al., 2022).

In our previous studies, all the electrophysiology, cardiorespiratory and pain studies were carried out in the rat, while the cell signalling assays and simulations were carried out using human receptors. Given the widespread availability and use of wild-type and mutant mice, we sought to determine whether the unique properties of BnOCPA were similar at the murine A_1_R in order to broaden the range of models with which to further study BnOCPA and adenosine signalling.

The A_1_Rs of human (hA_1_R), rat (rA_1_R), and mouse (mA_1_R) consist of 326 amino acids. The percentage of amino acid sequence identity of the A_1_R in the three species is human vs. rat 95 %, human vs. mouse 95 %, and rat vs. mouse 98 %, (Alnouri et al., 2015). Although there are only a small number of residues (six) that differ between the mA_1_R and rA_1_R, clear differences in the affinity and potency of ligands have previously been reported (Alnouri et al., 2015). For example, the agonist 2-Chloro-N^6^-cyclopentyladenosine (CCPA) has a much higher affinity at mouse receptors compared to rat, and PSB63 is a much more effective antagonist at rat receptors compared to murine receptors (Alnouri et al., 2015). These differences in ligand properties between the two receptors makes it plausible that there could indeed be differences in the properties of BnOCPA at the two receptors.

Here we show that the actions of BnOCPA are markedly different in the mouse vs the rat hippocampus. In the mouse, BnOCPA acts as an agonist at both pre- and postsynaptic A_1_Rs, to inhibit excitatory synaptic transmission and induce membrane hyperpolarisation, respectively, whereas in the rat, BnOCPA acts selectively at presynaptic receptors and does not induce membrane hyperpolarisation. Studies at recombinant rat, mouse and human A_1_Rs confirm BnOCPA’s preferential activation of Gob, but show no species differences in agonist binding, indicating that BnOCPA’s functional selectivity in the rat likely resides in G protein coupling.

## Results

### BnOCPA has a similar potency on the inhibition of synaptic transmission in mouse and rat hippocampus

We have previously shown that the selective A_1_R agonist BnOCPA selectively couples through Gob following the activation of the hA_1_R (Wall et al., 2022). This unique profile likely explains why BnOCPA has a differential action on pre- and postsynaptic hippocampal neurons in the rat hippocampal area CA1 (Wall et al., 2022). In the rat hippocampus, BnOCPA potently inhibited excitatory synaptic transmission (IC_50_ ∼65 nM) but even at high concentrations (tested up to 1 μM) did not hyperpolarise the membrane potential of CA1 hippocampal neurons, which, as evidenced by the use of Goa interfering peptides, predominately occurs via Goa activation (Wall et al., 2022). Here we have investigated if BnOCPA displays this differential behaviour in the mouse hippocampus.

We first tested if BnOCPA had a similar action at presynaptic mA_1_R in mouse compared to rat hippocampus. The effect of BnOCPA was measured on excitatory synaptic transmission (Figure 1). Using a concentration (120 nM), approximately twice the IC_50_ reported in rat hippocampus (Wall et al., 2022), we found that synaptic transmission was inhibited by 71.7 ± 2.5 % (*n* = 6 slices, Figure 1A). This is comparable with ∼76 % inhibition reported for this concentration of BnOCPA in the rat (measured from the concentration-response curve fit) (Wall et al., 2022). These effects of BnOCPA in the mouse hippocampus were slowly reversible on wash (Figure 1A) and could be reversed by the A_1_R antagonist 8CPT (2 μM; *n* = 3, Figure 1B, C). Thus, it appears that the potency of BnOCPA on presynaptic A_1_R in the hippocampus is comparable between mouse and rat.

**Figure 1.**
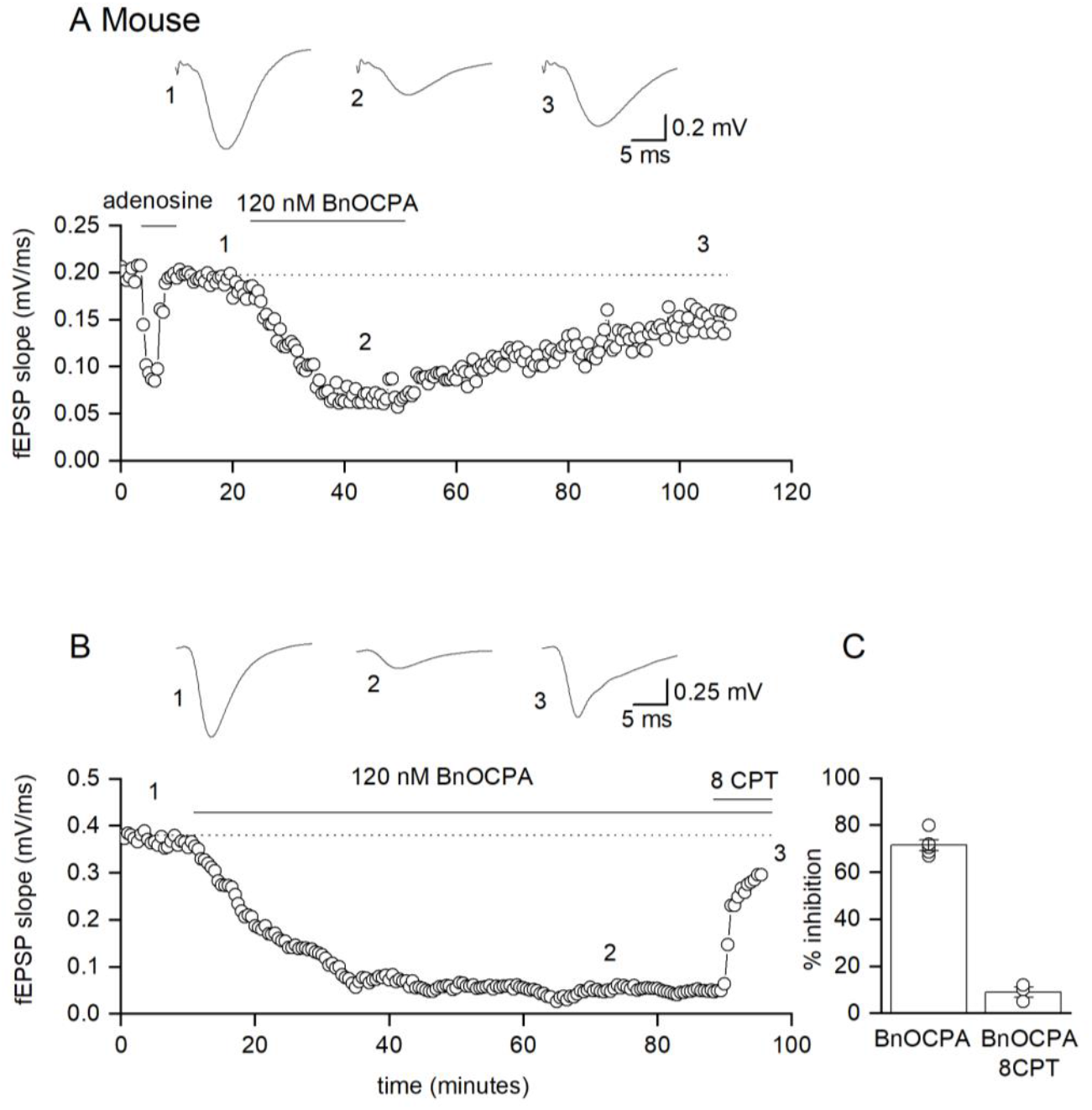
BnOCPA has approximately equal potency at mouse vs rat presynaptic A_1_Rs. A, fEPSP slope plotted against time for a single recording. Application of adenosine (50 μM) caused a reduction in fEPSP slope which rapidly recovered upon wash. Application of BnOCPA 120 nM reduced fEPSP slope by ∼70 %, an effect that was slowly reversible upon wash. Inset, average fEPSP waveforms in: 1) baseline, 2) BnOCPA 120 nM and 3) following wash. B, fEPSP slope plotted against time for a single recording. BnOCPA greatly reduced fEPSP slope, an effect greatly reduced by the A_1_ receptor antagonist 8CPT (2 μM). Inset, average fEPSP waveforms in 1) baseline, 2) BnOCPA 120 nM and 3) BnOCPA + 8CPT. C, Bar graph summarising inhibitory effects of BnOCPA (120 nM) and BnOCPA (120 nM) plus 8CPT (2 μM) on fEPSP slope. Points are data from individual recordings.

### BnOCPA hyperpolarises mouse hippocampal neurons, reduces their input resistance and firing rate

To examine the postsynaptic effects of BnOCPA on membrane potential, we applied BnOCPA to mouse hippocampal CA1 neurons in whole cell current clamp experiments, as we did previously in the rat (Wall et al., 2022). In contrast to previous observations with BnOCPA in rat CA1 neurones, application of 300 nM BnOCPA hyperpolarised the membrane potential of mouse CA1 neurons (mean hyperpolarisation 3.9 ± 0.7 mV, *n* = 6 slices, Figure 2A, C). This change in membrane potential could be partially reversed on prolonged wash and was accompanied by a fall in input resistance (mean reduction 19.4 ± 5.8 %, *n* = 6 slices, Figure 2B, D) and a large fall in firing rate (mean reduction 75.3 ± 6.5 %, as measured from the injection of a naturalistic current, see Methods, Figure 2E, F). These effects of BnOCPA were concentration-dependent, with reduced effects at lower concentrations (120 nM) and larger effects at higher concentrations (1 μM, Figure 2C, D and F). To confirm that the effects of BnOCPA were via the mA_1_R, we used the selective A_1_R antagonist 8-(4-Chlorophenylthio)adenosine (8CPT) (2 μM). Antagonism of A_1_Rs reversed the effects of BnOCPA on membrane potential (Figure 2G) and firing rate (Figure 2H). Thus, unlike in the rat, BnOCPA activates postsynaptic A_1_R in the mouse hippocampus at sub-micromolar concentrations, and in a manner consistent with the parent compound, the selective A_1_R agonist CPA.

**Figure 2.**
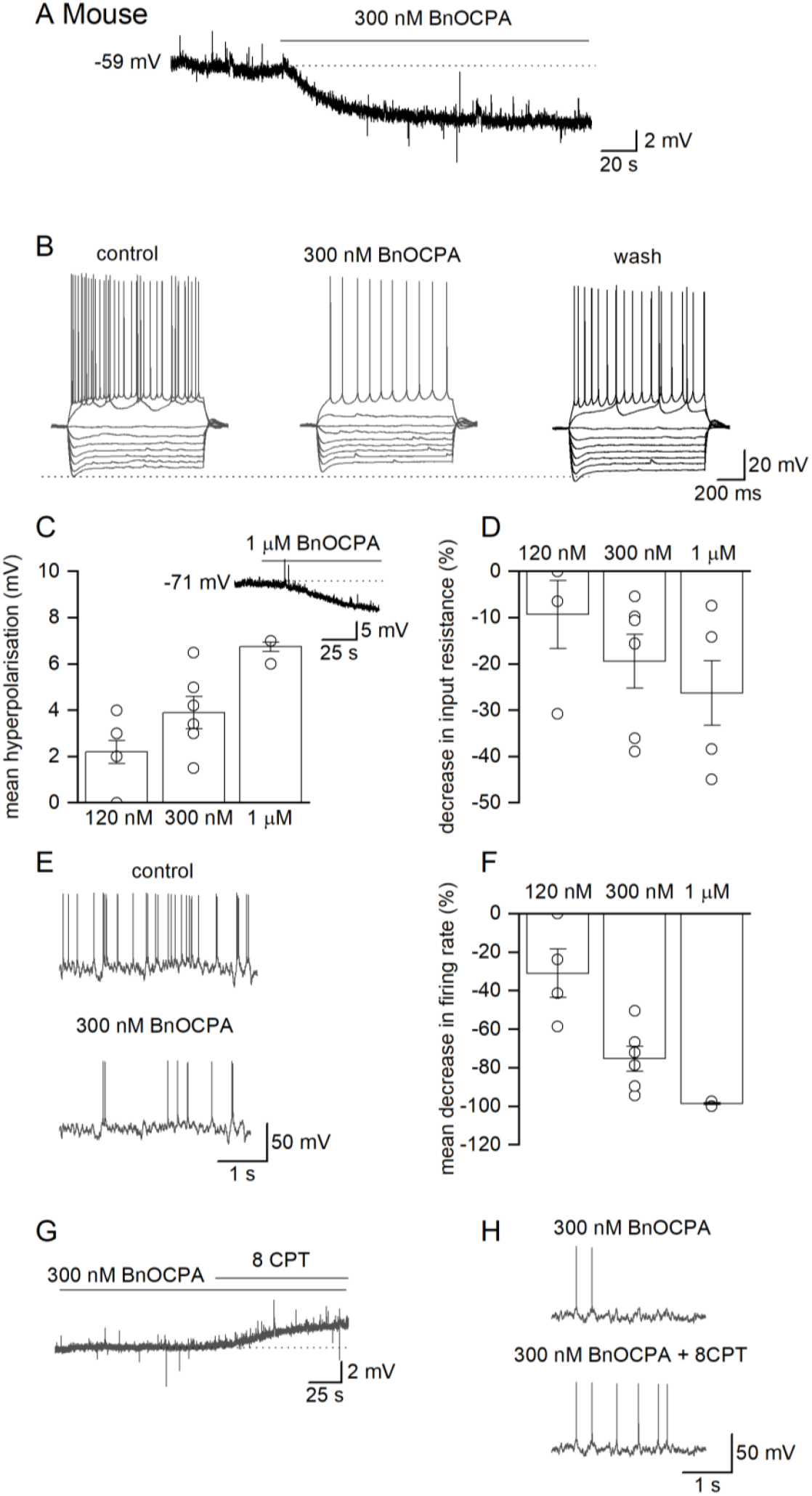
BnOCPA activates postsynaptic A_1_Rs on mouse hippocampal CA1 neurons. A, Membrane potential trace from a CA1 pyramidal cell in a mouse hippocampal slice. The application of 300 nM BnOCPA hyperpolarised the membrane potential by ∼ 3 mV (from – 59 to -62 mV). This effect was partially reversible upon wash (not shown). B, Current-voltage traces taken from the neuron in (A). The traces are aligned to the resting potential to illustrate the fall in input resistance in BnOCPA (measured before sag, dotted line). There is also an increase in firing threshold and a marked reduction in firing rate with positive current steps. For the current-voltage relations, the current steps started at -300 pA and were increased by 50 pA until steady firing occurred. C, Graph plotting mean hyperpolarisation of the membrane potential against concentration of BnOCPA. Inset, membrane potential trace showing large (∼7 mV) hyperpolarisation induced by 1 μM BnOCPA. D, Graph plotting the mean % reduction in input resistance against concentration of BnOCPA. E, Example of action potential firing induced by naturalistic current injection (see Methods) before BnOCPA application (top) and after BnOCPA (300 nM; bottom). Note the marked reduction in firing rate. F, Graph plotting the reduction (%) in firing rate against concentration of BnOCPA. Firing was abolished by 1 mM BnOCPA. For C, D and F the points are data points from individual experiments. G, Membrane potential trace from a CA1 pyramidal cell in a mouse hippocampal slice in the presence of 300 nM BnOCPA. Application of the A_1_R antagonist 8CPT (2 μM) reversed the hyperpolarisation caused by BnOCPA, resulting in depolarisation of the membrane potential by ∼2.5 mV. H, Example of action potential firing induced by naturalistic current injection (see Methods) during BnOCPA application (300 nM, top) and after application of 8CPT (2 μM, bottom), which restores action potential firing.

### Only high concentrations of BnOCPA produce a small hyperpolarisation of rat neurons

In interleaved experiments, using the same stock solution of BnOCPA as used in the mice recordings, we confirmed our previous data that BnOCPA does not hyperpolarise rat neurons at concentrations up to 1 μM. In 10 neurons (3 rats), application of 300 nM BnOCPA (∼5 times the IC_50_ for the inhibition of the fEPSP) had no appreciable effect on membrane potential (Figure 3A, B). Interestingly, there was no decrease in input resistance (as observed in mice) but instead the input resistance was slightly increased in some cells at this concentration (mean increase 9.9 ± 6.2 %, *n* = 10; Figure 3C, D). This increase in input resistance may reflect the reduction of tonic rA_1_R activation by endogenous adenosine as BnOCPA can prevent or reverse membrane hyperpolarisation caused by adenosine and CPA, and indeed acts like an antagonist when the receptor couples via Goa (Wall et al., 2022). However, this potential effect on tonic A_1_R activation had little effect on firing rate induced by current steps (Figure 3C) or injection of naturalistic membrane currents (Figure 3E).

**Figure 3.**
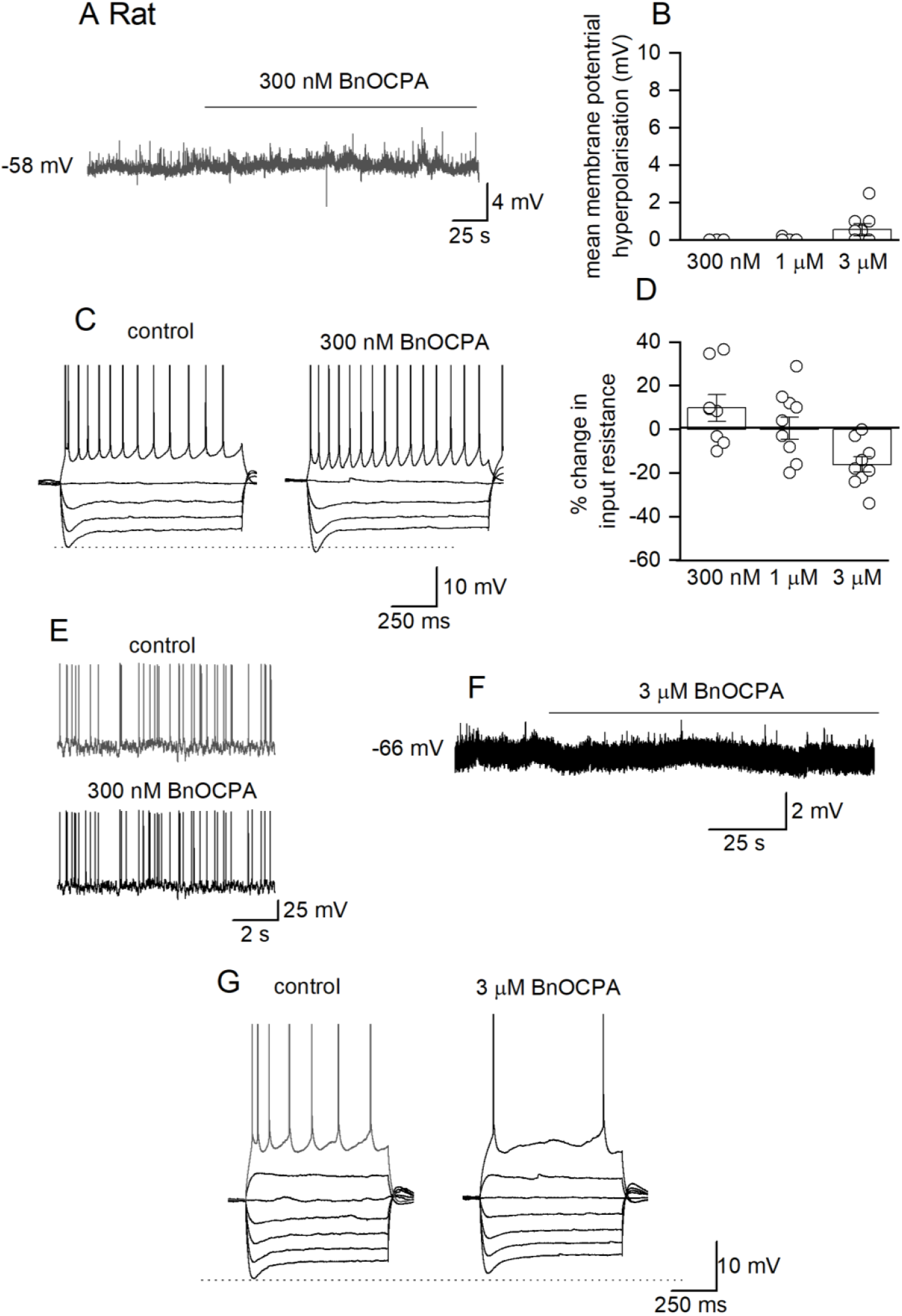
High concentrations of BnOCPA have little effect on rat CA1 hippocampal neurons. A, Membrane potential trace from a rat CA1 hippocampal neuron. Application of 300 nM BnOCPA had no effect. B, Graph plotting mean change in membrane potential (mV) against concentration of BnOCPA plotted on the same y axis scale as per the mouse (Figure 2C). Only at 3 μM does BnOCPA produce a small hyperpolarisation in some cells (mean ∼ 0.5 mV, compared to ∼ 6 mV for 1 μM in the mouse; Figure 2C). C, Current-voltage traces taken from the neuron in (A) before and after BnOCPA application. There was a small increase in input resistance (measured before the sag, as shown by the dotted line). Action potentials are truncated at 0 mV. D, Graph plotting mean change in input resistance against concentration of BnOCPA. The lower concentration of BnOCPA (300 nM) reflects the small increase in input resistance seen in Figure 3C. This changes to no net effect at 1 μM and reverses to a mean decrease of ∼15 % at 3 μM BnOCPA. E, Examples of action potential firing induced by naturalistic current injection (see Methods) before BnOCPA application (top) and after BnOCPA (300 nM) (bottom). There is a small increase in the firing rate in BnOCPA. F, Membrane potential trace from a rat CA1 hippocampal neuron. Application of 3 μM BnOCPA (∼50 times the IC_50_ for inhibiting synaptic transmission) had little effect. G, Example of a CA1 pyramidal neuron where 3 μM BnOCPA produced a small decrease in input resistance and a reduction in firing rate. Action potentials are truncated at 0 mV.

We have previously shown (Wall et al., 2022) and confirmed (Figure 3B) that at 1 μM BnOCPA (∼17 times the IC_50_ for the inhibition of the fEPSP) has no effect on membrane potential and had on average no consistent effect on input resistance (Figure 3D). Here we have increased the concentration of BnOCPA further to discover if it can hyperpolarise postsynaptic neurons. We found that at 3 μM (∼50 times the IC_50_ for the inhibition of the fEPSP) BnOCPA can hyperpolarise some CA1 neurons; the effect is still very small (mean change 0.5 mV; Figure 3B, F) and associated with a reduction in input resistance (Figure 3D) and a reduction in spike firing in response to currents steps (Figure 3G). This data supports our previous findings and shows that even at concentrations up to 3 μM BnOCPA has little effect on postsynaptic rA_1_Rs, in stark contrast to the observation made at mA_1_Rs.

### Assessing the binding affinity of BnOCPA at human A_1_R, rat A_1_R and mouse A_1_R

Having observed the differences in BnOCPA activity between the mouse and rat hippocampus, we compared the amino acid sequences between the human, rat and mouse A_1_Rs. There are only six residues that differ between rA_1_R and mA_1_R, three of which are located in the extracellular loop 2 (ECL2) and the other three are at the C-terminal (Figure 4). To examine whether the changes in ECL2 affected the binding affinities of A_1_R agonists, the affinities of NECA, adenosine, CPA and BnOCPA at hA_1_R, rA_1_R and mA_1_R were determined using a NanoBRET binding assay (Figure 5 and Table 1; (Preti et al., 2022)). In line with our previous findings (Deganutti et al., 2021), hA_1_R bound the fluorescent tracer CA200645 with a K_D_ of 67.4 ± 3.2 nM which was lower than both the rA_1_R (19.0 ± 2.5 nM) or mA_1_R (20.1 ± 1.5 nM). Among the four agonists explored, only NECA showed significantly different affinities between the species, higher at hA_1_R (pKi of 6.02 ± 0.03) when compared with rA_1_R (pKi of 5.76 ± 0.06) and mA_1_R (pKi of 5.72 ± 0.04). Importantly, BnOCPA displayed similar affinities at all three species (hA_1_R - pKi of 6.06 ± 0.13; rA_1_R - pKi of 6.16 ± 0.07 and mA_1_R - pKi of 6.21 ± 0.09). Therefore, it seems that although the differences in ECL2 might lead to different binding affinities for some agonists (Alnouri et al., 2015), they do not affect the affinity of BnOCPA at A_1_Rs across the three species tested.

**Figure 4.**
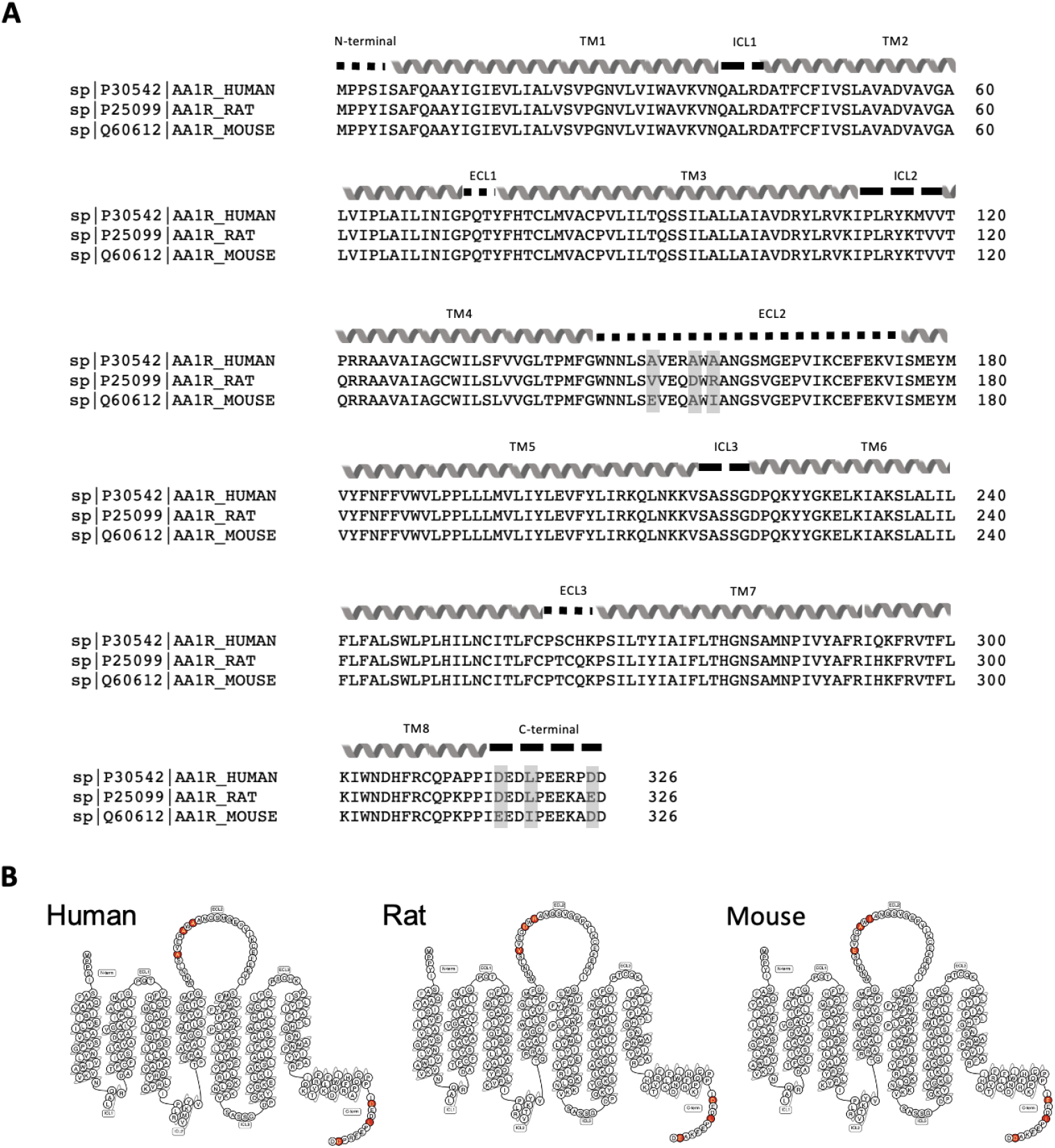
Amino acid sequence alignment of hA_1_R, rA_1_R and mA_1_R. A, The amino acid sequences were from Uniprot database (UniProt, 2020) and the multiple sequence alignment was performed in Clustal Omega programme (Sievers et al., 2011). There are 6 out of 326 residues differing between human, rat and mouse A_1_Rs and are all highlighted with grey boxes. They are: A/V/E151, A/D/A155 and A/R/I157 located at the ECL2, and D/D/E316, L/L/I319 and D/E/D326 at the C-terminal of hA_1_R/rA_1_R/mA_1_R respectively. B, Snake plot of hA_1_R, rA_1_R and mA_1_R with the critical residues highlighted with red circles. The plot was generated with the webtools in GPCRdb.org (Isberg et al., 2013).

**Figure 5.**
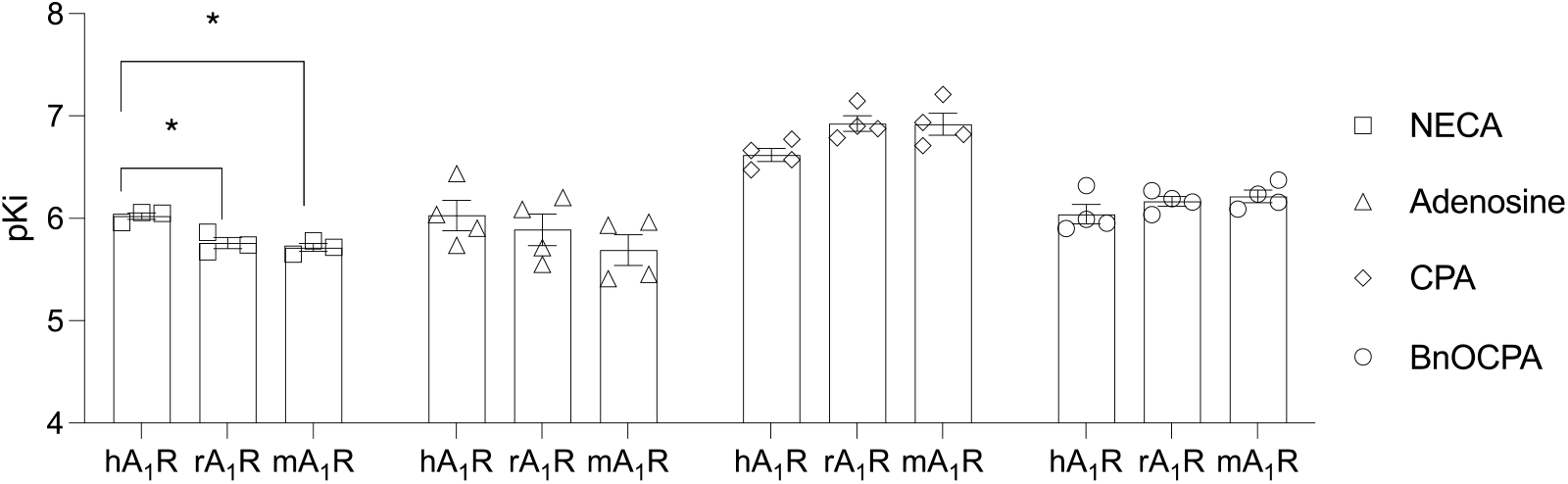
BnOCPA has similar binding affinities at hA_1_R, rA_1_R and mA_1_R. Increasing concentrations of unlabelled AR ligands together with 70 nM or 20 nM of CA200645 were added to HEK293 cells transiently transfected with Nluc-hA_1_R or Nluc-rA_1_R/Nluc-mA_1_R. The pKi values were determined using Cheng-Prusoff equation and plotted as mean ± SEM of at least three independent experiments performed in duplicates. A One-way ANOVA with a Tukey’s multiple comparison test was used to test for statistically significant (\**p* < 0.05) differences between interspecies agonist affinity. Only NECA showed significant differences between the hA_1_R and the rA_1_R, and the hA_1_R and the mA_1_R. See Table 1 For actual p values.

**Table 1.**
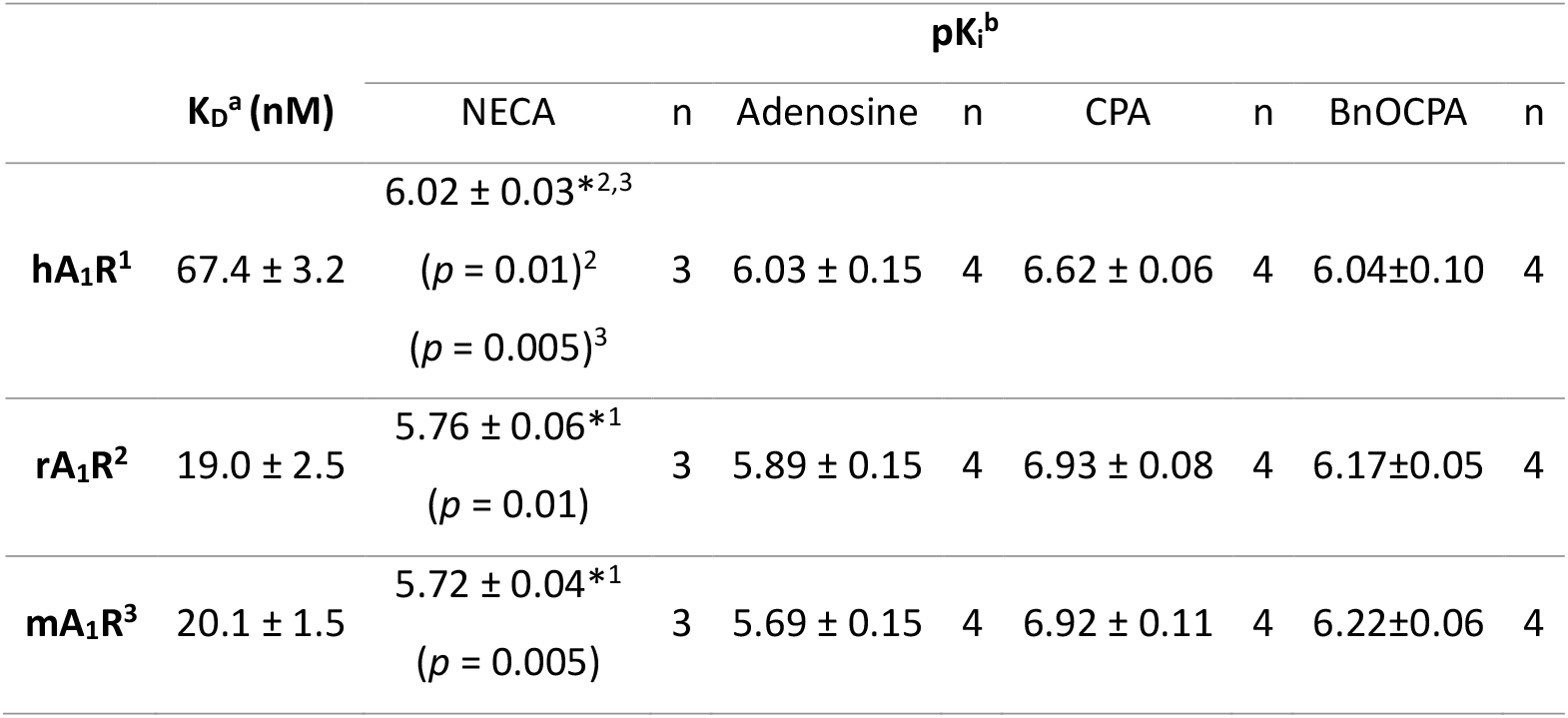
NanoBRET Saturation-binding and competition-binding assay at hA_1_R, rA_1_R and mA_1_R. NanoBRET Saturation and competition assay were used to determine the ^a^equilibrium constant (KD) of CA200645 and the compound affinity (pKi), respectively. HEK293T cells transiently transfected with Nluc-hA_1_R, Nluc-rA_1_R or Nluc-mA_1_R were used to calculate the corresponding K_D_ and pK_i_ at the hA_1_R, rA_1_R and mA_1_R, respectively. Data are presented as mean ± SEM of at least three independent experiments conducted in duplicate. In the competition assay, 70 nM or 20 nM of CA200645 were used to compete with unlabelled agonists at hA_1_R, rA_1_R or mA_1_R. The baseline-corrected BRET ratio values at 10 minutes post-stimulation were used to determine the binding affinity of compounds. A One-way ANOVA with Tukey’s multiple comparison tests were used to compare the statistical significance (\**p* < 0.05) of the interspecies difference of the agonist affinity. *^x^ denotes p < 0.05 when compared with x=1, hA_1_R; x=2, rA_1_R and x=3, mA_1_R and exact *p* values are provided for relevant comparisons.

### The selective activation of Gob over Goa by BnOCPA-induced stimulation of A_1_R is conserved across different species

We previously reported the selective activation of Gob over Goa by BnOCPA in functional cAMP and TRUPATH assays, which may explain its differential actions on pre- and postsynaptic A_1_Rs in rat (Wall et al., 2022). To investigate whether BnOCPA displays this discrimination between Goa and Gob at A_1_R from different species, we again turned to utilising the BRET-based TRUPATH G protein activation assay (Olsen et al., 2020). We first examined the Goa and Gob activation induced by BnOCPA and three other AR agonists (NECA, CPA and adenosine), stimulating hA_1_R, rA_1_R or mA_1_R. In line with our previous observations, BnOCPA showed significantly higher potency at activating Gob over Goa at both hA_1_R and rA_1_R (Figure 6A and 6B). Moreover, this selectivity for Gob was observed in the case of mA_1_R. Importantly, NECA, adenosine and CPA failed to display significantly higher potency at activating Gob over Goa at all the three A_1_Rs (Figure 6C). Thus, the lack of discrimination between pre- and postsynaptic A_1_Rs in mice was not due to the loss of bias in activating Gob over Goa by BnOCPA.

**Figure 6.**
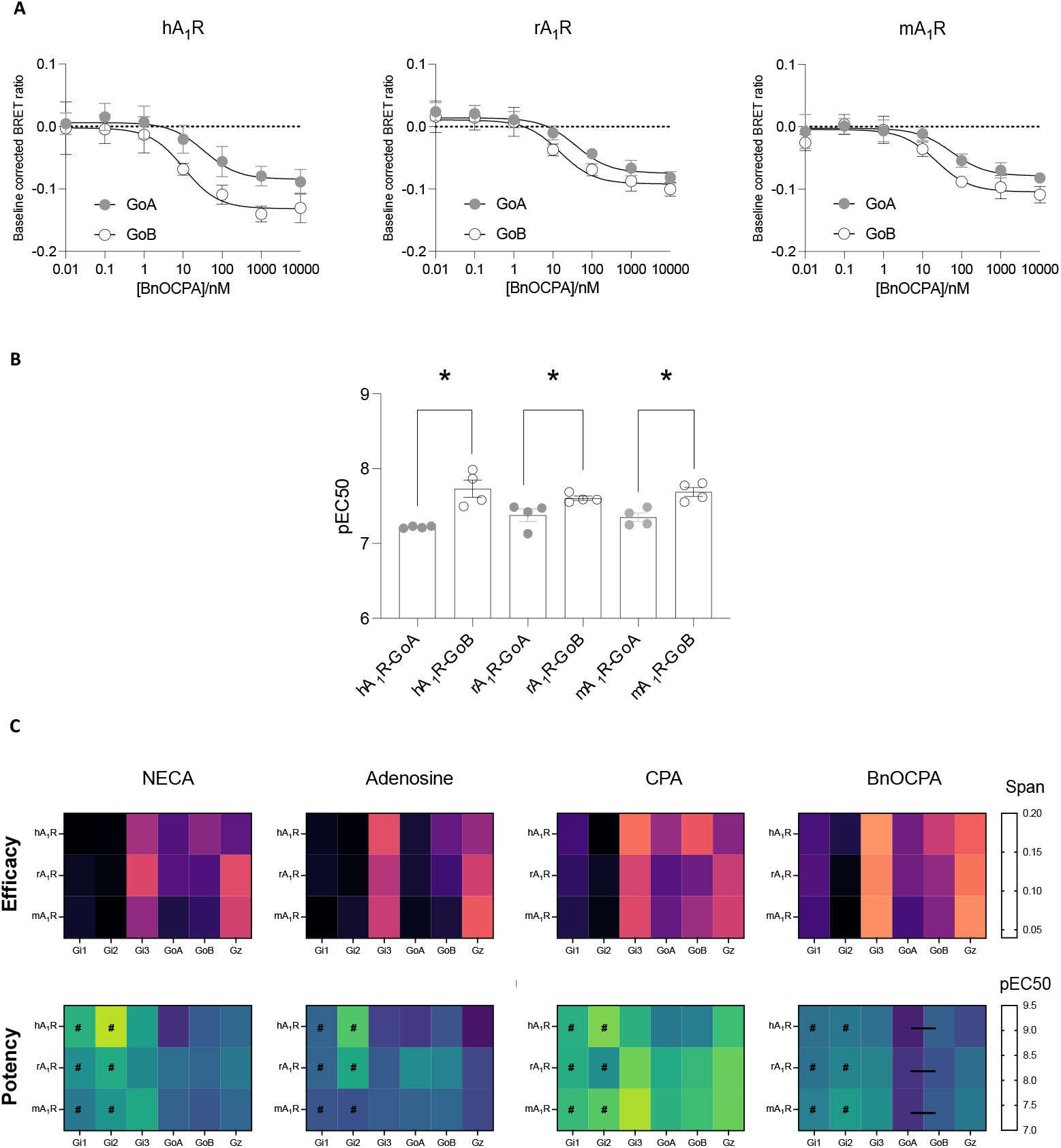
BnOCPA has uniquely selective A_1_R activation of Gob over Goa across all three species while NECA, adenosine and CPA do not show similar discrimination between Goa and Gob. A, BnOCPA-stimulated activation of Goa and Gob at hA_1_R, rA_1_R and mA_1_R determined in the TRUPATH G protein assay. The BRET2 values at 11 minutes poststimulation were used to generate concentration-response curves. B, The comparison of potency (pEC_50_) in Goa vs Gob activation. BnOCPA selectively activated Gob over Goa at hA_1_R (*p* = 0.005), rA_1_R (*p* = 0.042) and mA_1_R (*p* = 0.007). C, Heatmaps summarising the span and pEC_50_ for individual Gi/o proteins. Among all the four AR agonists tested, only BnOCPA displayed greater activation of Gob over Goa (in potency) at rA_1_R, and this selective activation was observed with the A_1_Rs from all three species. # denotes the pEC_50_ obtained from a response with very small span (< 0.05) which has less reliability. Data are represented as mean ± SEM in A, B, and mean in C from at least three independent experiments conducted in duplicate. Unpaired t-tests were used to calculate the statistical significance between Goa and Gob activation level (\**p* < 0.05, with actual *p* values given above).

Apart from Goa and Gob, there are other non-sensory Gi/o proteins also expressed in CNS (Sjöstedt et al., 2020) so they might contribute to the differential action of A_1_Rs. Therefore, we expanded our exploration by investigating the agonist-induced activation of all six non-sensory G_i/o_ proteins at hA_1_R, rA_1_R or mA_1_R with four different AR agonists (Figure 6C and Table 2). All three A_1_Rs were able to activate each of the six non-sensory Gi/o proteins although the efficacy (as determined by the span of the response) for G_i1_ and G_i2_ were barely above the baseline of the assay; thus any calculation of potency is unreliable (denoted as # in Figure 6C). We did notice robust responses at G_i3_ and G_z_ when stimulated with all four agonists, although caution is needed in comparing the efficacy ranges due to differences in the Gβ*γ* combinations used in the TRUPATH assay. We observed that CPA was able to activate the rA_1_R with a significantly higher potency than that of the hA_1_R at G_i3_. Furthermore, when we compared the potencies of BnOCPA at the three A_1_Rs, we observed that it displayed a significantly higher potency for the rA_1_R compared to hA_1_R at activating Gi3, and higher potency for the mA_1_R compared to hA_1_R at activating Gz, although there was no significant difference between the rA_1_R and mA_1_R. Thus, in terms of agonist binding and Gα-coupling we were unable to identify any mechanism for the observed differences in pre- and postsynaptic activity of BnOCPA at the rA_1_R and mA_1_R.

**Table 2.** Gi/o protein activation level at hA_1_R, rA_1_R and mA_1_R stimulated by four AR agonists. Potency and span of NECA/Adenosine/CPA/BnOCPA-stimulated activation of six non-sensory Gi/o proteins at hA_1_R, rA_1_R and mA_1_R determined in the TRUPATH G protein activation assay. # denotes the pEC_50_ obtained from a response with very small span (*p* < 0.05) which has less reliability. Data are presented as mean ± SEM from at least three independent experiments conducted in duplicate. One-way ANOVA with Tukey’s multiple comparisons tests were performed to calculate significant difference between hA_1_R^1^, rA_1_R^2^ and mA_1_R^3^ at activating each Gi/o protein. *^x^ denotes p < 0.05 when compared with x=1, hA_1_R; x=2, rA_1_R and x=3, mA_1_R and exact *p* values are provided for relevant comparisons.

| <u>Potency</u> |  |  |  |  |  |  |  |  |  |  |  |  |  |
| --- | --- | --- | --- | --- | --- | --- | --- | --- | --- | --- | --- | --- | --- |
|  | **pEC50** | **G<sub>i1</sub>** | **n** | **G<sub>i2</sub>** | **n** | **G<sub>i3</sub>** | **n** | **G<sub>oA</sub>** | **n** | **G<sub>oB</sub>** | **n** | **G<sub>z</sub>** | **n** |
| NECA | hA<sub>1</sub>R<sup>1</sup> | 8.59±0.24<sup>#</sup> | 4 | 9.25±0.13<sup>#</sup> | 3 | 8.40±0.15 | 5 | 7.33±0.14 | 3 | 7.71±0.13 | 5 | 7.82±0.24 | 5 |
|  | rA<sub>1</sub>R<sup>2</sup> | 8.23±0.20<sup>#</sup> | 4 | 8.51±0.25<sup>#</sup> | 4 | 8.16±0.09 | 4 | 7.89±0.14 | 3 | 7.80±0.10 | 4 | 7.99±0.11 | 4 |
|  | mA<sub>1</sub>R<sup>3</sup> | 8.00±0.04<sup>#</sup> | 3 | 8.30±0.31<sup>#</sup> | 3 | 8.48±0.17 | 3 | 7.67±0.34 | 3 | 7.67±0.23 | 3 | 7.81±0.29 | 3 |
| Adenosine | hA<sub>1</sub>R<sup>1</sup> | 7.79±0.30<sup>#</sup> | 5 | 8.82±0.60<sup>#</sup> | 5 | 7.49±0.15 | 6 | 7.40±0.12 | 5 | 7.64±0.12 | 5 | 7.12±0.09 | 5 |
|  | rA<sub>1</sub>R<sup>2</sup> | 7.79±0.26<sup>#</sup> | 4 | 8.48±9.76<sup>#</sup> | 4 | 7.69±0.12 | 4 | 8.19±0.37 | 3 | 8.10±0.35 | 4 | 7.51±0.24 | 4 |
|  | mA<sub>1</sub>R<sup>3</sup> | 7.65±0.15<sup>#</sup> | 3 | 7.62±0.23<sup>#</sup> | 3 | 7.72±0.23 | 3 | 7.78±0.23 | 3 | 7.82±0.25 | 3 | 7.53±0.09 | 3 |
| CPA | hA<sub>1</sub>R<sup>1</sup> | 8.56±0.13<sup>#</sup> | 4 | 9.01±0.11<sup>#</sup> | 3 | 8.60±0.09<sup>*3</sup><br>(*p*=0.005) | 4 | 8.00±0.15 | 4 | 8.05±0.07 | 4 | 8.71±0.03 | 4 |
|  | rA<sub>1</sub>R<sup>2</sup> | 8.55±0.24<sup>#</sup> | 4 | 8.21±0.67<sup>#</sup> | 4 | 8.96±0.12 | 4 | 8.51±0.06 | 3 | 8.66±0.30 | 4 | 8.92±0.05 | 4 |
|  | mA<sub>1</sub>R<sup>3</sup> | 8.68±0.31<sup>#</sup> | 5 | 8.84±0.37<sup>#</sup> | 5 | 9.20±0.06<sup>*1</sup><br>(*p*=0.005) | 3 | 8.60±0.23 | 5 | 8.71±0.22 | 5 | 8.93±0.18 | 5 |
| BnOCPA | hA<sub>1</sub>R<sup>1</sup> | 7.90±0.15<sup>#</sup> | 6 | 8.00±0.10<sup>#</sup> | 5 | 7.89±0.03<sup>*2</sup><br>(*p*=0.02) | 6 | 7.22±0.07 | 4 | 7.73±0.12 | 4 | 7.56±0.12<sup>*3</sup><br>(*p*=0.04) | 6 |
|  | rA<sub>1</sub>R<sup>2</sup> | 7.94±0.07<sup>#</sup> | 6 | 8.11±0.07<sup>#</sup> | 6 | 8.12±0.07<sup>*1</sup><br>(*p*=0.02) | 6 | 7.38±0.08 | 4 | 7.61±0.03 | 4 | 7.88±0.08 | 6 |
|  | mA<sub>1</sub>R<sup>3</sup> | 8.07±0.02<sup>#</sup> | 4 | 8.37±0.25<sup>#</sup> | 4 | 8.22±0.15 | 4 | 7.35±0.06 | 4 | 7.69±0.06 | 4 | 8.01±0.14<sup>*1</sup><br>(*p*=0.04) | 4 |

**Table 2 Gi/o protein activation level at hA<sub>1</sub>R, rA<sub>1</sub>R and mA<sub>1</sub>R stimulated by four AR agonists**
|  | **Span** | **G<sub>i1</sub>** | **n** | **G<sub>i2</sub>** | **n** | **G<sub>i3</sub>** | **n** | **G<sub>oA</sub>** | **n** | **G<sub>oB</sub>** | **n** | **G<sub>z</sub>** | **n** |
| --- | --- | --- | --- | --- | --- | --- | --- | --- | --- | --- | --- | --- | --- |
| NECA | hA<sub>1</sub>R | 0.043±0.005 | 4 | 0.044±0.003 | 5 | 0.114±0.008 | 5 | 0.083±0.005 | 5 | 0.104±0.011 | 5 | 0.087±0.005 | 5 |
|  | rA<sub>1</sub>R | 0.052±0.009 | 4 | 0.046±0.006 | 4 | 0.135±0.002 | 3 | 0.083±0.009 | 3 | 0.082±0.011 | 4 | 0.138±0.014 | 4 |
|  | mA<sub>1</sub>R | 0.055±0.001 | 3 | 0.043±0.009 | 3 | 0.106±0.012 | 3 | 0.063±0.007 | 3 | 0.070±0.004 | 3 | 0.131±0.015 | 3 |
| Adenosine | hA<sub>1</sub>R | 0.047±0.006 | 5 | 0.040±0.004 | 5 | 0.137±0.007 | 6 | 0.054±0.003 | 6 | 0.086±0.004 | 6 | 0.105±0.009 | 5 |
|  | rA<sub>1</sub>R | 0.049±0.007 | 4 | 0.042±0.004 | 4 | 0.119±0.017 | 4 | 0.053±0.005 | 3 | 0.072±0.006 | 4 | 0.129±0.005 | 4 |
|  | mA<sub>1</sub>R | 0.038±0.004 | 3 | 0.051±0.004 | 3 | 0.126±0.012 | 3 | 0.045±0.009 | 3 | 0.056±0.008 | 3 | 0.141±0.006 | 3 |
| CPA | hA<sub>1</sub>R | 0.074±0.014 | 4 | 0.039±0.003 | 4 | 0.154±0.007 | 4 | 0.109±0.005 | 3 | 0.144±0.008 | 4 | 0.102±0.006 | 4 |
|  | rA<sub>1</sub>R | 0.067±0.006 | 4 | 0.049±0.002 | 4 | 0.133±0.009 | 4 | 0.081±0.003 | 4 | 0.090±0.001 | 4 | 0.128±0.006 | 4 |
|  | mA<sub>1</sub>R | 0.060±0.006 | 5 | 0.044±0.005 | 5 | 0.137±0.012 | 5 | 0.088±0.005 | 5 | 0.111±0.004 | 5 | 0.131±0.011 | 5 |
| BnOCPA | hA<sub>1</sub>R | 0.077±0.005 | 6 | 0.060±0.009 | 6 | 0.164±0.008 | 6 | 0.092±0.004 | 6 | 0.126±0.005 | 4 | 0.145±0.010 | 6 |
|  | rA<sub>1</sub>R | 0.080±0.004 | 6 | 0.045±0.005 | 6 | 0.159±0.011 | 6 | 0.092±0.003 | 6 | 0.104±0.004 | 6 | 0.153±0.008 | 6 |
|  | mA<sub>1</sub>R | 0.074±0.005 | 4 | 0.039±0.003 | 4 | 0.163±0.009 | 4 | 0.076±0.004 | 4 | 0.099±0.002 | 4 | 0.161±0.006 | 4 |

## Discussion

There is an appreciable difference in the actions of BnOCPA in the mouse hippocampus compared to the rat, with the unique functional properties of BnOCPA, the selective activation of pre-vs postsynaptic A_1_Rs, not present in the mouse. One possible reason for the different effects of BnOCPA in rats and mice could be differential coupling between the A_1_R, G proteins and the K^+^ channels that mediate the hyperpolarisation. In the rat, we have shown that the A_1_R couples through Goa to activate K^+^ channels leading to membrane hyperpolarisation (Wall et al., 2022). It is possible that A_1_R in mouse hippocampal neurons instead couple via Gob to activate K^+^ channels. This, however, seems unlikely as expression of Goa is high in the postnatal brain of both mice and rats, with little Gob expression (Rouot et al., 1992; Kolasa et al., 2000).

We speculated that perhaps a more plausible explanation was that differences in A_1_R sequence between rats and mice changes the binding of BnOCPA leading to differential G protein coupling. However, using NanoBRET-binding assays we were able to demonstrate that the differences in the ECL2 between rat and mouse did not change the binding affinity of BnOCPA at A_1_Rs. Moreover, these residue differences between rA_1_R and mA_1_R did not affect the selectivity for activating Gob over Goa elicited by BnOCPA based on the results from TRUPATH G protein activation assay.

However, there are several limitations in the TRUPATH G protein activation assay which might result in failing to explain the different effects observed in rat and mouse pre- and postsynaptic A_1_Rs from the perspective of GPCR-G protein interactions. First, all the G protein subunit components in this assay are based on human G proteins. The sequences of G proteins vary across species so even with equal potency in activating the same G protein, the G protein from different species might have different effects in activating the downstream effectors. For example, the residues at the H3.05 position in rat Goa and Gob are Ser and Phe, respectively, whereas they are both Ser in mouse Goa and Gob (Figure 7). This different residue in rat Gob might lead to the amplification of the difference between Goa and Gob potency in rat while this amplification is lost in mice. Second, the combinations of heterotrimeric Gαβγ protein are fixed in the TRUPATH assay for optimising the output level; both Goa and Gob are used in combination with Gβ3 and Gγ8 in this system (Olsen et al., 2020). Notwithstanding the interspecies sequence differences of Gβ3 and Gγ8 subunits, there are five different Gβ protein family members and twelve Gγ protein family members (Hurowitz et al., 2000), leading to a wide range of possible combinations. It is possible that different combinations of Gβ and Gγ subunits are recruited in rat/mouse pre- and postsynaptic membranes, either because of the differences in the expression levels of Gβ and Gγ subunits, or due to the different residues at the C-terminal of the rA_1_R and mA_1_R. The mechanisms underlying the different actions of BnOCPA on rat and mouse pre- and post-synaptic A_1_Rs remain to be elucidated in future experiments by examining the interspecies differences between the various G proteins.

**Figure 7.**
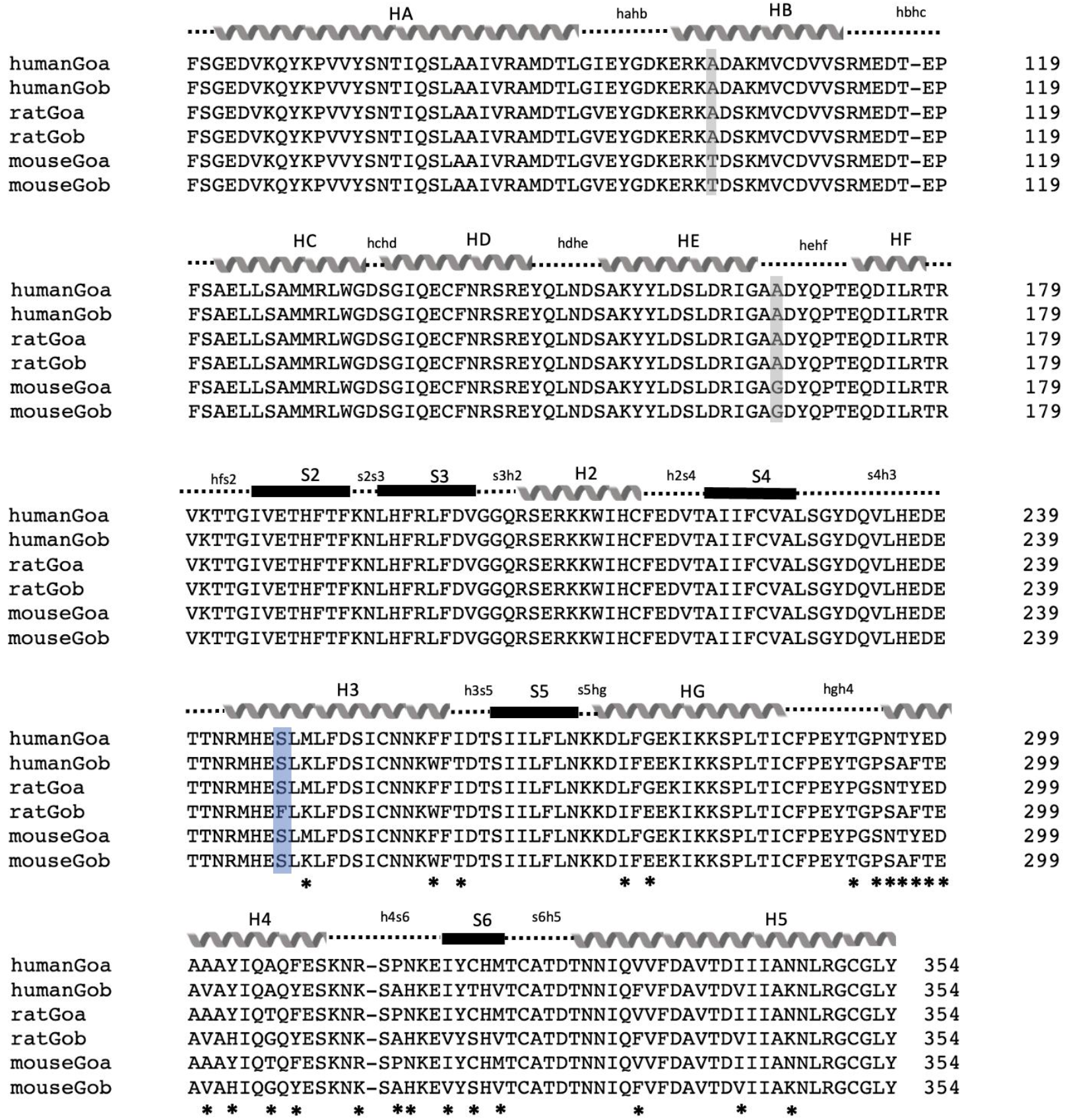
Amino acid sequence alignment of human, rat and mouse Goa/Gob. Amino acid sequences were from Tsukamoto et al., (Tsukamoto et al., 1991) and the multiple sequence alignment was performed in Clustal Omega programme (Sievers et al., 2011). There are only three positions that differ between rat/mouse Goa/Gob. Mouse Goa/Gob have unique residues at the positions highlighted in grey box when compared with human or rat Goa/Gob. At the position H3.05 (numbered in common Gα naming system), rat Gob has unique phenylalanine instead of the widely conserved serine. Black stars indicate the positions that are non-conserved between Goa and Gob, but these differences are present in all three species.

## Acknowledgements

We gratefully acknowledge the support of the BBSRC (BB/W1014831/1, G.L., and BB/M01116X/1, Ph.D. studentship to I.D.P.), the MRC (2270402, iCASE PhD Studentship to C.L.M.) and The Swiss National Science Foundation (Swiss National Science Foundation PP00P2_123536 and PP00P2_146321 M.L.). X.H. is funded by a China Scholarship Council Cambridge International Scholarship, and E.H. by a Race Against Dementia Fellowship (ARUK-RADF2021A009). G.L. is a Royal Society Industry Fellow (INF\R2\212001).

## Methods

### Drugs and solutions

For the electrophysiology studies, drugs were made up as stock solutions (1-10 mM, stored at -20 °C) and then diluted in aCSF or saline on the day of use. BnOCPA ((2*R*,3*R*,4*S*,5*R*)-2-(6-{[(1*R*,2*R*)-2-benzyloxycyclopentyl]amino}-9*H*-purin-9-yl)-5-(hydroxymethyl)oxolane-3,4-diol) was synthesised as described previously (Knight et al., 2016) and dissolved in dimethyl-sulfoxide (DMSO, 0.01% final concentration). Adenosine, 8-CPT (8-cyclopentyltheophylline) were purchased from Sigma-Aldrich (Poole, Dorset, UK). For the cell signalling assays, adenosine, NECA, CPA and DPCPX (8-cyclopentyl-1,3-dipropyl-7H-purine2,6-dione) were purchased from Tocris Bioscience (Wiltshire, UK) and dissolved in DMSO as 10mM stocks, stored at -20°C until use. CA200645 was purchased from HelloBio (Bristol, UK), dissolved in DMSO as 100 μM stock and stored at -20°C.

### Approvals

All experiments involving animals were conducted with the knowledge and approval of the University of Warwick Animal Welfare and Ethical Review Board, and in accordance with the U.K. Animals (Scientific Procedures) Act (1986) and the EU Directive 2010/63/EU.

### Preparation of hippocampal slices

Sagittal slices of hippocampus (300-400 μm) were prepared from male Sprague Dawley rats and C57BL/6J mice, at postnatal days 12-20. Rats and mice were kept on a 12-hour light-dark cycle with slices made 90 minutes after entering the light cycle. In accordance with the U.K. Animals (Scientific Procedures) Act (1986), rats and mice were humanely killed by cervical dislocation and then decapitated. The brain was removed, cut down the midline and the two sides of the brain stuck down to a metal base plate using cyanoacrylate glue. Slices were cut along the midline with a Microm HM 650V microslicer in cold (2-4°C), high Mg^2+^, low Ca^2+^ artificial cerebrospinal fluid (aCSF), composed of (mM): 127 NaCl, 1.9 KCl, 8 MgCl_2_, 0.5 CaCl_2_, 1.2 KH_2_PO_4_, 26 NaHCO_3_, 10 D-glucose (pH 7.4 when bubbled with 95% O_2_ and 5% CO_2_, 300 mOSM). Slices were stored at 34°C for 1-6 hours in aCSF (1 mM MgCl_2_, 2 mM CaCl_2_) before use.

### Extracellular recording

A slice was transferred to the recording chamber, submerged in aCSF and perfused at 4-6 ml·min^-1^ (32 ± 0.5°C). The slice was placed on a grid allowing perfusion above and below the tissue and all tubing was gastight (to prevent loss of oxygen). An aCSF-filled glass microelectrode was placed within stratum radiatum in area CA1 and recordings were made using a differential model 3000 amplifier (AM systems, WA USA). Field excitatory postsynaptic potentials (fEPSPs) were evoked with an isolated pulse stimulator model 2100 (AM Systems, WA). For fEPSPs a 10-20 minute baseline was recorded at a stimulus intensity that gave 40-50 % of the maximal response before drug application. Signals were acquired at 10 kHz, filtered at 3 kHz and digitised on-line (10 kHz) with a Micro CED (Mark 2) interface controlled by Spike software (Vs 6.1, Cambridge Electronic Design, Cambridge UK) For fEPSP slope, a 1 ms linear region after the fibre volley was measured.

### Whole-cell patch clamp recording from hippocampal pyramidal cells

A slice was transferred to the recording chamber and perfused at 3 ml·min^-1^ with aCSF at 32 ± 0.5°C. Slices were visualized using IR-DIC optics with an Olympus BX151W microscope (Scientifica) and a CCD camera (Hitachi). Whole-cell recordings were made from pyramidal cells in area CA1 of the hippocampus using patch pipettes (5–10 MΩ) manufactured from thick walled glass (Harvard Apparatus, Edenbridge UK) and containing (mM): potassium gluconate 135, NaCl 7, HEPES 10, EGTA 0.5, phosphocreatine 10, MgATP 2, NaGTP 0.3 and biocytin 1 mg ml^−1^ (290 mOSM, pH 7.2). Recordings were obtained using an Axon Multiclamp 700B amplifier (Molecular Devices, USA) and digitised at 20 KHz. Data acquisition and analysis was performed using Pclamp 10 (Molecular Devices, USA).

Current−voltage relationships were constructed by injecting step currents from either –200 or -300 pA incrementing by either 50 or 100 pA (1 s) until a regular firing pattern was induced. A plot of step current against voltage response around the resting potential was used to measure the input resistance (gradient of the fitted line, before the voltage sag). To induce firing, a current wave form, designed to provoke naturalistic fluctuating voltages, was constructed using the summed numerical output of two Ornstein–Uhlenbeck processes (Uhlenbeck and Ornstein, 1930) with time constants τfast = 3 ms and τslow = 10 ms. This current wave form, which mimics the background postsynaptic activity resulting from activation of AMPA and GABA_A_ receptor channels, is injected into cells and the resulting voltage recorded (a fluctuating, naturalistic trace of 40s in duration). The firing rate was measured from voltage traces evoked by injecting a current wave form of the same gain for all recordings (firing rate ∼2–3 Hz). APs were detected by a manually set threshold and the interval between APs measured.

### Constructs

Both sigNluc-hA_1_R and sigNluc-rA_1_R in pcDNA3.1+ were kindly gifted by Prof Stephen Hill and Dr Steve Briddon (University of Nottingham). The pcDNA5/FRT-hA_1_R and pcDNA5/FRT-rA_1_R were generated by Dr Kerry Barkan. The original construct of mA_1_R was purchased from Bio-Techne Ltd. (Minneapolis, US) and then cloned into sigNluc-pcDNA3.1+ vector and pcDNA5/FRT vector (Thermo Fisher Scientific) respectively. The Gαi/o-RLuc8, Gβ and Gγ-GFP2 constructs were purchased as part of the TRUPATH kit from Addgene.

### Cell culture and transfection

HEK293T cells were gifted by Jürgen Müller (University of Warwick) and were maintained in DMEM/F-12 GlutaMAX™ media (ThermoFisher, UK), with 10 % FBS (Sigma-Aldrich, Poole, Dorset, UK) and 1 % AA (Sigma, UK). HEK293T cells were plated in a 6-well plate and grown overnight. The seeded cells were then transfected with a total 2 μg of DNA using polyethylenimine 25 kDa (PEI, Polysciences Inc., Germany) at a 6:1 ratio of PEI to DNA, diluted in 150 mM NaCl. All cells were maintained at 37 °C with 5 % CO_2_, in a humidified atmosphere.

### NanoBRET binding assay

In both the NanoBRET saturation and competition binding assay, 2 μg of Nluc-hA_1_R, Nluc-rA_1_R or Nluc-mA_1_R was transfected into the cells seeded in 6-well plate. After 24 h, the cells were seeded onto white 96-well plates (Greiner, UK) at a density of 10,000 cells/well and grown overnight. The medium was discarded and replaced with 100 μl phosphate buffered saline (PBS) (ThermoFisher, UK) containing 0.1 % Bovine Serum Albumin (BSA, Sigma-Aldrich, Poole, Dorset, UK) and 0.05 μM Nano-Glo Luciferase substrate (Promega, UK). After five-minute incubation, different concentrations (0.1 nM-1000 nM) of CA200645 were added to the saturation assay while different concentrations (1 nM-100 μM) of unlabelled AR agonists together with CA200645 (70 nM for hA_1_R and 20 nM for rA_1_R/mA_1_R) were added in the competition assay. Non-specific binding was determined with the addition of 10 μM DPCPX, an unlabelled antagonist. Light emissions at 460 nm (Nluc) and >610 nm (fluorescent ligand) were measured using Mithras LB940 for 30 min. The raw BRET ratio was calculated by dividing the emission at 610 nM with the 450 nM emission and then baseline-corrected. The baseline-corrected BRET ratio at 10 min poststimulation was used to determine the KD of CA200645 as well as the pKi of AR agonists at hA_1_R, rA_1_R and mA_1_R, respectively.

### TRUPATH G protein activation assay

The seeded cells in 6-well plate were transfected with hA_1_R/rA_1_R/mA_1_R, Gα-RLuc8, Gβ, Gγ-GFP2 and pcDNA3.1 with the ratio of 1:1:1:1:1 using 400 ng per construct together. The combinations of Gα Gβ and Gγ used were the same as the previous literature suggested (Olsen et al., 2020). After 24 h transfection, cells were seeded onto poly-L-lysine (PLL)-coated white 96-well plates (Greiner, UK) at a density of 50,000 cells/well in a complete DMEM/F12 medium and grown overnight. On the day of assay, the culture media was discarded and replaced with 80 μl assay buffer (1× Hank’s balanced salt solution (HBSS) with calcium, supplemented with 20 mM HEPES and 0.1 % BSA, pH 7.4). 10 μl of coelenterazine 400a (Nanolight technology, USA) was added to the assay buffer for a final concentration of 5 μM. The plates were then incubated in the dark for 5 minutes, followed by the addition of 10 μl AR agonists (0.01 nM – 10 μM). BRET signal was recorded every 60 seconds for 20 minutes on a Pherastar plate reader and the BRET ratio was calculated as the ratio of light emission from GFP2 (515 nm) over Rluc8 (400 nm). Net BRET ratio was baseline-corrected and the value at 11 minute poststimulation was used to generate the concentration response curves for data analysis.

### Data and statistical analysis

All the cell-based assay data were analysed with Prism 9.4.1 (Graphpad, San Diego, CA). The dissociation constant (KD) of CA200645 at hA_1_R, rA_1_R and mA_1_R was determined by fitting the BRET ratio dose-response curve with the “one-site total model” in saturation binding built into Prism. The affinity constant (pKi) of the AR agonists in NanoBRET ligand binding competition assay was determined by fitting the baseline-corrected BRET ratio response curve with the “one-site Ki model” in competitive binding based on Cheng and Prusoff equation (Cheng and Prusoff, 1973) with both the concentration (RadioligandNM) and Kd (HotKDNM) values of the ‘Hot ligand’ CA200645 set to 70 nM and 20 nM for hA_1_R and rA_1_R/mA_1_R respectively. One-way ANOVA with Turkey’s multiple comparisons was used to calculate the significant difference (*p < 0.05) between the binding affinity of AR ligands at A_1_R from different species. In the TRUPATH G protein activation assay, baseline-corrected concentration-response curves were fitted with a three-parameter logistics equation to determine the response potency (pEC_50_), minimal response (E_min_), maximal response (E_max_) and span (E_max_-E_min_). Unpaired t-tests were used to calculate the significant difference (* p < 0.05) in the potency of Goa and Gob at different A_1_Rs elicited by the AR agonists tested. One-way ANOVA with Tukey’s multiple comparisons tests were performed to calculate the significant difference between hA_1_R, rA_1_R and mA_1_R at activating each Gi/o protein. Statistical significance for the pharmacological data was determined as described in (Curtis et al., 2018).

